# The role of “spillover” in antibiotic resistance

**DOI:** 10.1101/536714

**Authors:** Scott W. Olesen, Marc Lipsitch, Yonatan H. Grad

**Author notes:** Correspondence: Yonatan Grad, Harvard T. H. Chan School of Public Health, 665 Huntington Ave, Building 1, Room 715, Boston, Massachusetts 02115.

## Abstract

Antibiotic use is a key driver of antibiotic resistance. Understanding the quantitative association between antibiotic use and resulting resistance is important for predicting future rates of antibiotic resistance and for designing antibiotic stewardship policy. However, the use-resistance association is complicated by “spillover”, in which one population’s level of antibiotic use affects another population’s level of resistance via the transmission of bacteria between those populations. Spillover is known to have effects at the level of families and hospitals, but it is unclear if spillover is relevant at larger scales. We used mathematical modeling and analysis of observational data to address this question. First, we used dynamical models of antibiotic resistance to predict the effects of spillover. Whereas populations completely isolated from one another do not experience any spillover, we found that if even 1% of interactions are between populations, then spillover may have large consequences: the effect of a change in antibiotic use in one population on antibiotic resistance in that population could be reduced by as much as 50%. Then, we quantified spillover in observational antibiotic use and resistance data from US states and European countries for 3 pathogen-antibiotic combinations, finding that increased interactions between populations were associated with smaller differences in antibiotic resistance between those populations. Thus, spillover may have an important impact at the level of states and countries, which has ramifications for predicting the future of antibiotic resistance, designing antibiotic resistance stewardship policy, and interpreting stewardship interventions.

## INTRODUCTION

Antibiotic resistance is a major threat to public health (1). Outpatient antibiotic use, which accounts for approximately 80% of human antibiotic use (2,3), is considered a principal driver of antibiotic resistance in the community (4). Understanding the relationship between use and resistance is important because it allows accurate predictions of the future of antibiotic resistance and goal-oriented antibiotic stewardship policy. The use-resistance association has been previously characterized in many ecological studies at the level of US states (5–7) and European countries (8,9). However, antibiotic resistance is a complex, temporally dynamic phenomenon (10–13), and many factors complicate the use-resistance association, making what should be an “obvious” connection sometimes difficult to identify and quantify (11). Even when detected, observed use-resistance associations are sometimes weaker than might be expected (14). One factor that could account for the difficulty in detecting use-resistance associations in ecological studies and for the apparent weakness of such associations is “spillover”.

“Spillover” is a consequence of the fact that antibiotic-resistant and -susceptible bacteria can be transmitted from person to person. Thus, one person’s risk of an antibiotic resistant infection depends on their own antibiotic use (15,16) as well as the rates of antibiotic use among their contacts (17). For example, one person’s use of antibiotics increases the risk of an antibiotic resistant infection among their family members (18–21). As another example, hospitalized patients with no recent antibiotic use can have a higher risk of resistance than people in the community with high antibiotic use (22) because antibiotic use and resistance in other hospitalized patients are high.

Spillover is important for three reasons. First, it means antibiotic resistance is not merely a localized problem. It is well-understood that new resistance determinants can emerge in one geography and spread globally (23,24), but the role of spillover in determining the levels of resistance in a given locale is not well-quantified. To what degree, for example, can one US state expect that its antibiotic resistance levels are due to antibiotic use within its borders, rather than in surrounding states? Second, spillover makes it difficult to design antibiotic stewardship interventions and understand their results. For example, if antibiotic use in one hospital changes, resistance might not change as expected because of spillover, from the community or other hospitals into that hospital’s patients. Finally, spillover makes it difficult to interpret the results of controlled antibiotic interventions, such as the effect of mass drug administration on antibiotic resistance (25,26), when the intervention and control populations are not wholly epidemiologically separate.

The effect of spillover should scale with the amount of interaction between populations. If two populations do not interact at all, then antibiotic use in one population cannot affect resistance in the other. However, if two populations liberally exchange bacteria, then the rates of antibiotic resistance in the two populations will be very similar, regardless of whether their rates of antibiotic use differ greatly.

The effects of spillover also depend on population sizes. For example, two large populations will have most interactions within themselves, rather than between each other. Spillover should therefore be most pronounced when considering small populations and become less important for large populations. As mentioned above, a single individual’s risk of resistance is modulated by antibiotic use in their family or in their healthcare facility. Spillover is also observed at the level of hospitals, as the level of resistance in one hospital appears to be affected by resistance levels in nearby hospitals as well as by antibiotic use rates in the surrounding communities (27–29). Presumably, when examining ever larger populations, such as US Census tracts (30), US states, or European countries, the effect of spillover will become less important. However, the relationship between population size and spillover effects is not well understood.

We hypothesized that US states and European countries, which are large populations with relatively independent public health policies, may be subject to substantially lower levels of antibiotic resistance spillover than family-or hospital-sized populations. This hypothesis, if true, would mean that individual states or countries could act as independent “laboratories” of antibiotic use and resistance. If not, it means that outpatient antibiotic resistance policy must be national or international in order to achieve its full effect. To evaluate this hypothesis, we first use mathematical models of antibiotic use and resistance to make quantitative predictions about the effect of spillover between populations as a function of their amount of mutual interaction. Then, we search for signals of spillover in observational data of antibiotic use and resistance in US states and European countries.

## METHODS

### Dynamical model of antibiotic resistance

To examine how interactions between populations could theoretically affect the association between antibiotic use and resistance, we used the within-host neutrality (WHN) mathematical model presented by Davies *et al*. (31) and described in the Supplemental Methods. Briefly, the model predicts the prevalence *ρ* of antibiotic resistance that results from an antibiotic use rate *τ* in a single, well-mixed population. To verify that conclusions drawn from the WHN model are not specific to the model structure, we also repeated all analyses with the “D-types” model of use and resistance (32). We selected these two models because they demonstrate coexistence between sensitive and resistant strains at equilibrium over a wide parameter space. Parameter values and simulation methodology for both models are in the Supplemental Methods. In the simulations, antibiotic use is measured as monthly treatments per capita and resistance as the proportion of colonized hosts carrying resistant strains.

To conceptually frame and clarify the question of spillover, we simulated an antibiotic stewardship intervention experiment using a structured host population approach inspired by Blanquart *et al*. (33). We considered pairs of an intervention population with antibiotic use rate *τ*_int_ and a control population with use rate *τ*_cont_. To determine how spillover affects the intervention’s measured outcome, we modulated the proportion *ε* of each population’s contacts that are in the other population. For *ε* = 0%, the populations are completely separate. For *ε* = 50%, contacts across populations are just as likely as contacts within populations (Supplemental Methods). We varied *ε* between 0% and 50%, and we varied the difference in use *Δτ* = *τ*_cont_ – *τ*_int_ between 0 and 0.15 treatments per person per month while keeping the average use 0.5 × (*τ*_cont_ + *τ*_int_) fixed at 0.125, reflecting the range of antibiotic use rates in the original model presentations.

### Observational data

We examined antibiotic use and resistance for 3 pathogen-antibiotic combinations: *S. pneumoniae* and macrolides, *S. pneumoniae* and β-lactams, and *Escherichia coli* and quinolones. We considered these 3 combinations because they are the subject of many modeling (31,32) and empirical studies (5,15).

Observational data were drawn from 3 sources. First, we used MarketScan (34) and ResistanceOpen (35) as previously described (7). The MarketScan data includes outpatient pharmacy antibiotic prescription claims for 62 million unique people during 2011-2014. ResistanceOpen includes antibiotic resistance data collected during 2012-2015 from 230 hospitals, laboratories, and surveillance units in 44 states. Second, we used the QuintilesIMS Xponent database (36) and the US Centers for Disease Control and Prevention’s (CDC) National Healthcare Safety Network (NHSN) (37). The Xponent data includes state-level data on US quinolone use during 2011-2014. NHSN includes state-level data on quinolone resistance among *E. coli* catheter-associated urinary tract infections during 2011-2014. Third, we used the European Center for Disease Prevention and Control’s (ECDC) ESAC-Net antimicrobial consumption database (38) and EARS-Net Surveillance Atlas of Infectious Disease (39) for 2011-2015. The ESAC-Net data includes country-level outpatient antibiotic use data provided by WHO and Ministries of Health from member countries. The EARS-Net data includes country-level resistance data. In the observational data, we quantified antibiotic use as yearly treatments per capita and resistance as the proportion of collected isolates that were non-susceptible. Further details about preparation of these data sources and their availability are in the Supplemental Methods.

We excluded the *S. pneumoniae* resistance to β-lactams in US states from the analysis because, in previous work using the same primary datasets, the point estimate for the use-resistance relationship was negative (14).

### Use-resistance relationships by populations’ adjacency

To test the theoretical prediction that the same difference in antibiotic use will be associated with smaller differences in antibiotic resistance when two populations (US states or European countries) have stronger interactions, we tested whether the use-resistance association is weaker in adjacent pairs of populations, which presumably have more cross-population contacts, compared to non-adjacent populations. Two populations were considered adjacent if they share a land or river border (40–42).

We quantified the use-resistance association as the percentage point difference (i.e., absolute risk difference) in resistance (proportion of non-susceptible isolates) divided by the difference in antibiotic use. We summarized use-resistance associations among adjacent pairs and non-adjacent pairs of populations using the median value. Because use-resistance associations between pairs of populations are correlated, we used the jackknife method to compute confidence intervals on the difference in medians between groups. We used the Mann-Whitney *U* test to compute statistical significance. Because our theoretical results suggested the ratio of percentage points difference in resistance divided by difference in antibiotic use rates was a predictable function of the degree of population mixing, we considered only this functional form for the use-resistance association.

### Use-resistance associations by interactions

Because adjacency might be too coarse measure of populations’ interactions to detect spillover, we performed a similar analysis as above, but predicting the use-resistance association between populations using transportation data. For US states, we used inter-county commuting statistics from the US Census (43). For European countries, we used inter-country passenger flight data from Eurostat (44). Rather than trying to infer a precise mathematical relationship between transportation statistics and epidemiological contacts, we used a nonparametric approach: we assumed that pairs of populations with relatively little inter-population transportation also have relatively few inter-population contacts, but we infer only the rank ordering of inter-population contacts, not their magnitudes. Specifically, we first converted the matrix of the number of counts (workers in the commuting data and passengers in the flight data) from each population to every other population into the proportion of counts moving from one population to another (i.e., divided each row by its sum), then symmetrized the resulting matrix (taking the elementwise average of the matrix and its transpose), and finally converted the resulting values into ranks. We assumed that intra-population interactions outnumber inter-population interactions and so set diagonal entries, which represent within-population interactions, to the highest rank. We measured the association between ranked interactions and use-resistance associations using the nonparametric Spearman’s correlation, computed confidence intervals using the jackknife method, and tested for statistical significance using the Mantel test with 999 permutations. To quantify the effect of spillover, we compared the median use-resistance associations among the top and bottom decile of ranked interactions.

Simulations and observational analyses were made using R (version 3.6.0) (45). The Mantel test used the *vegan* package (46). Multiple hypotheses were accounted for using Benjamini-Hochberg false discovery rate.

## RESULTS

In simulations of two populations, representing an intervention and control group, interactions between the two populations attenuated the effect of the intervention (Figure 1). With increasing interaction strength, the same intervention, that is, the same difference in antibiotic use between the populations, was associated with a smaller difference in antibiotic resistance. The difference in resistance between populations increases with the difference in antibiotic use (Figure 1d), but the use-resistance association, measured as the ratio of the difference in resistance to the difference in use, depends strongly on the interaction strength (Figure 1e). Thus, spillover between populations attenuates the measured use-resistance association.

**Figure 1.**
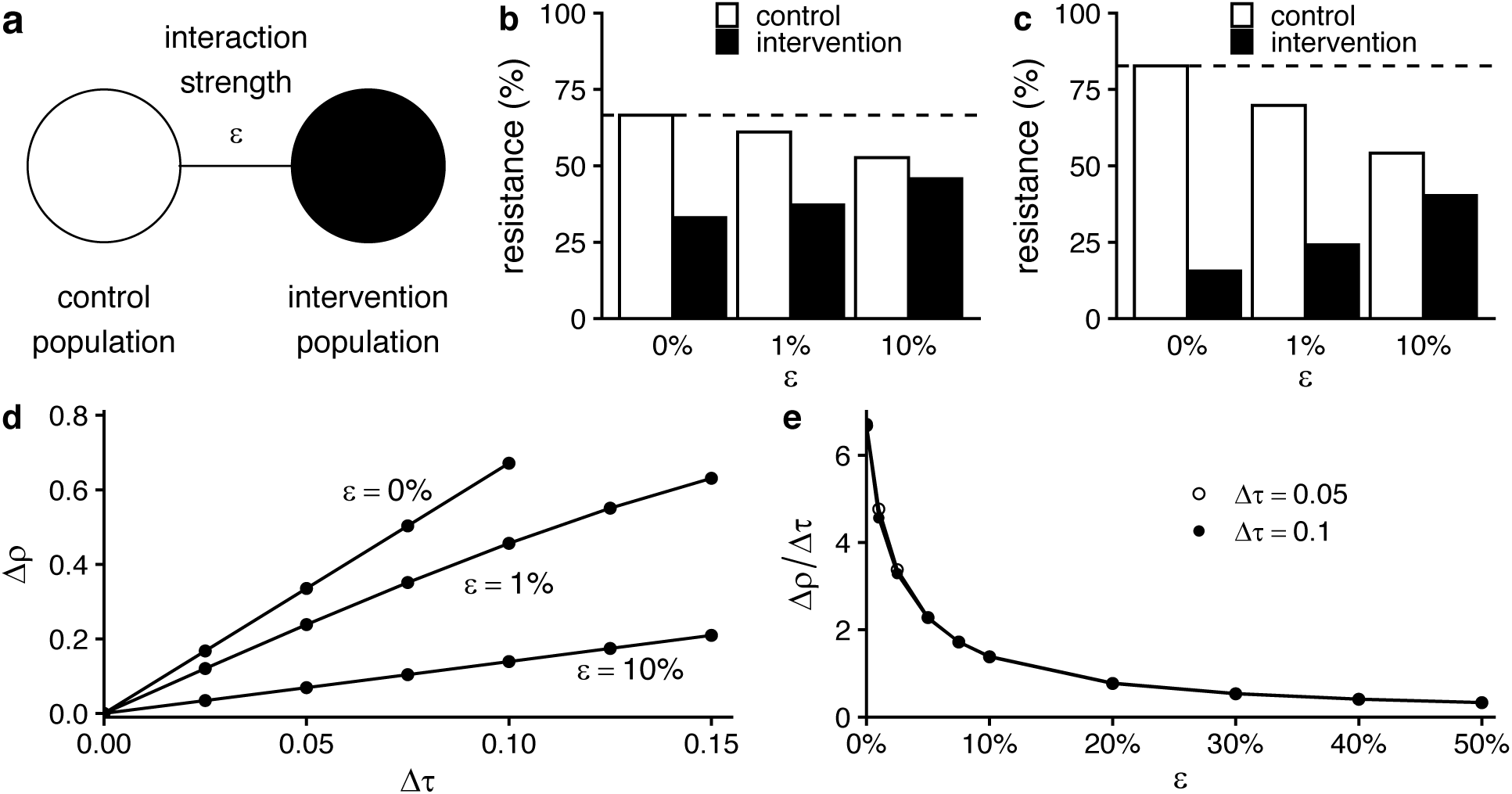
Interactions between populations attenuate the effect of interventions. (*a*) Schematic of the 2-population model. (*b*) Results of simulations of the 2-population WHN model for a modest intervention (difference in antibiotic use between populations Δ*τ* = 0.05 monthly treatments per capita; average of control and intervention treatment rates 0.125). As interaction strength (*ε*, horizontal axis) increases, the difference in antibiotic resistance between the two populations decreases. Dotted line shows resistance level in populations before the intervention. (*c*) The same pattern holds for a stronger intervention (Δ*τ* = 0.1, same average treatment rate). (*d*) In general, the difference in resistance between populations (Δ*ρ*, vertical axis) increases with the difference in antibiotic use (Δ*τ*, horizontal axis). (*e*) However, in the WHN model, the use-resistance relationship (Δ*ρ*/Δ*τ*, vertical axis) depends mostly on the interaction strength *ε* and is mostly independent of the difference in antibiotic use Δ*τ*.

The precise relationship between *ε*, the proportion of each population’s contacts that are in the other population, and the attenuation of the use-resistance association depended on the choice of mathematical model (Supplemental Table 1, Supplemental Figure 1). For *ε* = 1%, the use-resistance declined by approximately 30% in the WHN model and more than 60% in the “D-types” model. In other words, the models predict that as few as 1% of contacts need to be across populations, rather than within populations, to cause the observed effect of an antibiotic stewardship intervention to shrink by one-third, or even half.

To test whether spillover is important at the scale of US states or European countries, we measured use-resistance associations between pairs of populations in 6 combinations of pathogen species, antibiotic class, and data source (Figure 2). We reasoned that if spillover is relevant at these scales, then pairs of states or countries with stronger interactions would have detectably weaker use-resistance associations.

**Figure 2.**
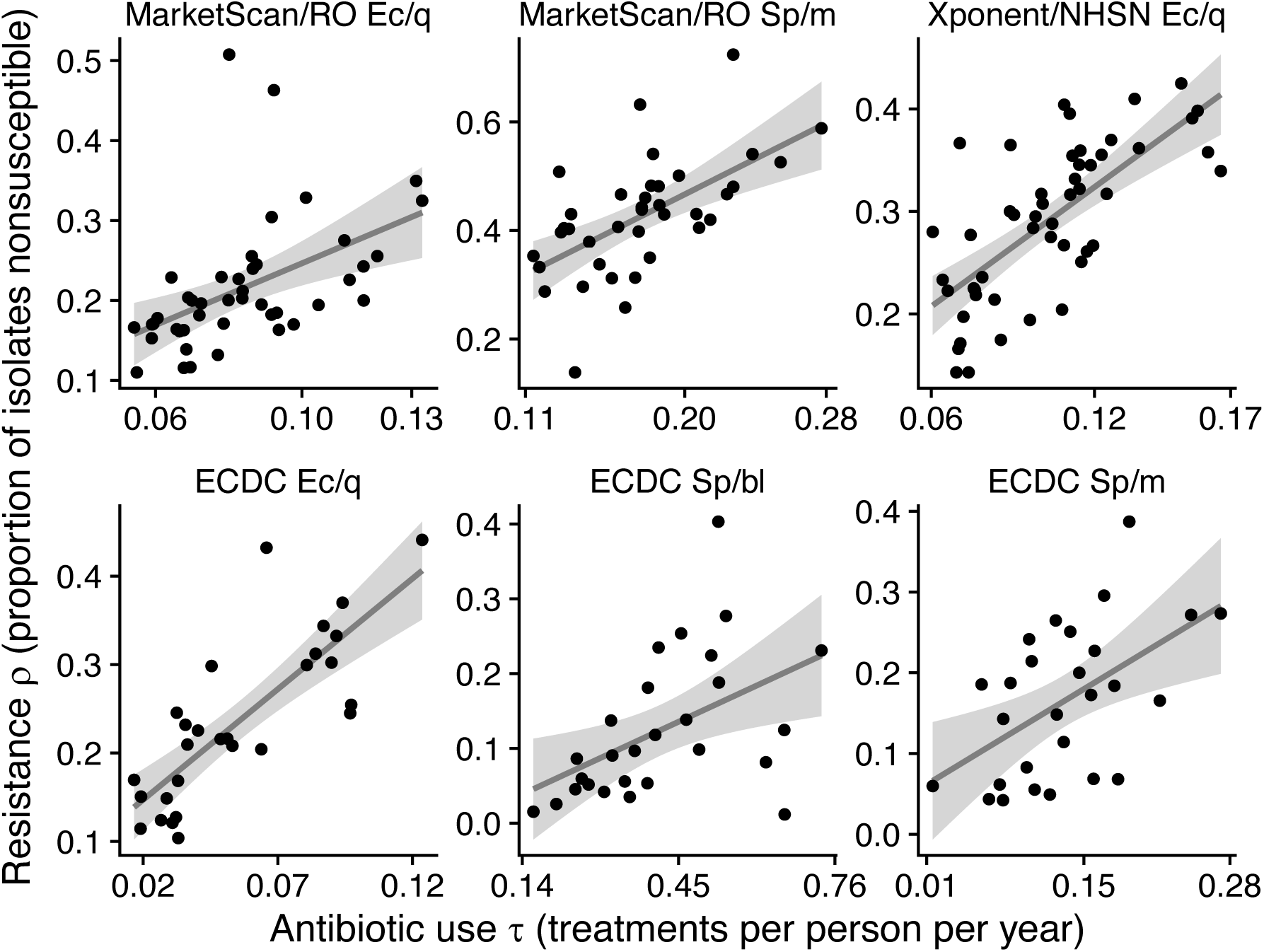
Use-resistance relationships across US states and European countries. Each point represents antibiotic use and resistance in a US state (top row) or European country (bottom row). Lines show simple linear regression best fit. Gray areas show 95% confidence interval. Ec/q: *E. coli* and quinolones. Sp/m: *S. pneunomiae* and macrolides. Sp/bl: *S. pneumoniae* and β-lactams. RO: ResistanceOpen. ECDC: European CDC.

We first tested whether pairs of physically adjacent populations (e.g., Massachusetts and Connecticut) had weaker use-resistance associations than non-adjacent populations (e.g., Massachusetts and Alaska). In 5 of 6 pathogen/antibiotic/dataset combinations, the median use-resistance association was smaller among adjacent populations than among non-adjacent populations (Figure 3), but in no case was the difference statistically significant after multiple hypothesis correction (Supplemental Table 2). Point estimates of the relative difference in median use-resistance associations between adjacent populations were 18% to 50% weaker than between non-adjacent populations (excepting *S. pneumoniae* with β-lactams, which was an outlier), consistent with the theoretical modeling results showing a 50% reduction in the use-resistance association for populations with approximately 1% of interactions across populations (Supplemental Table 1).

**Figure 3.**
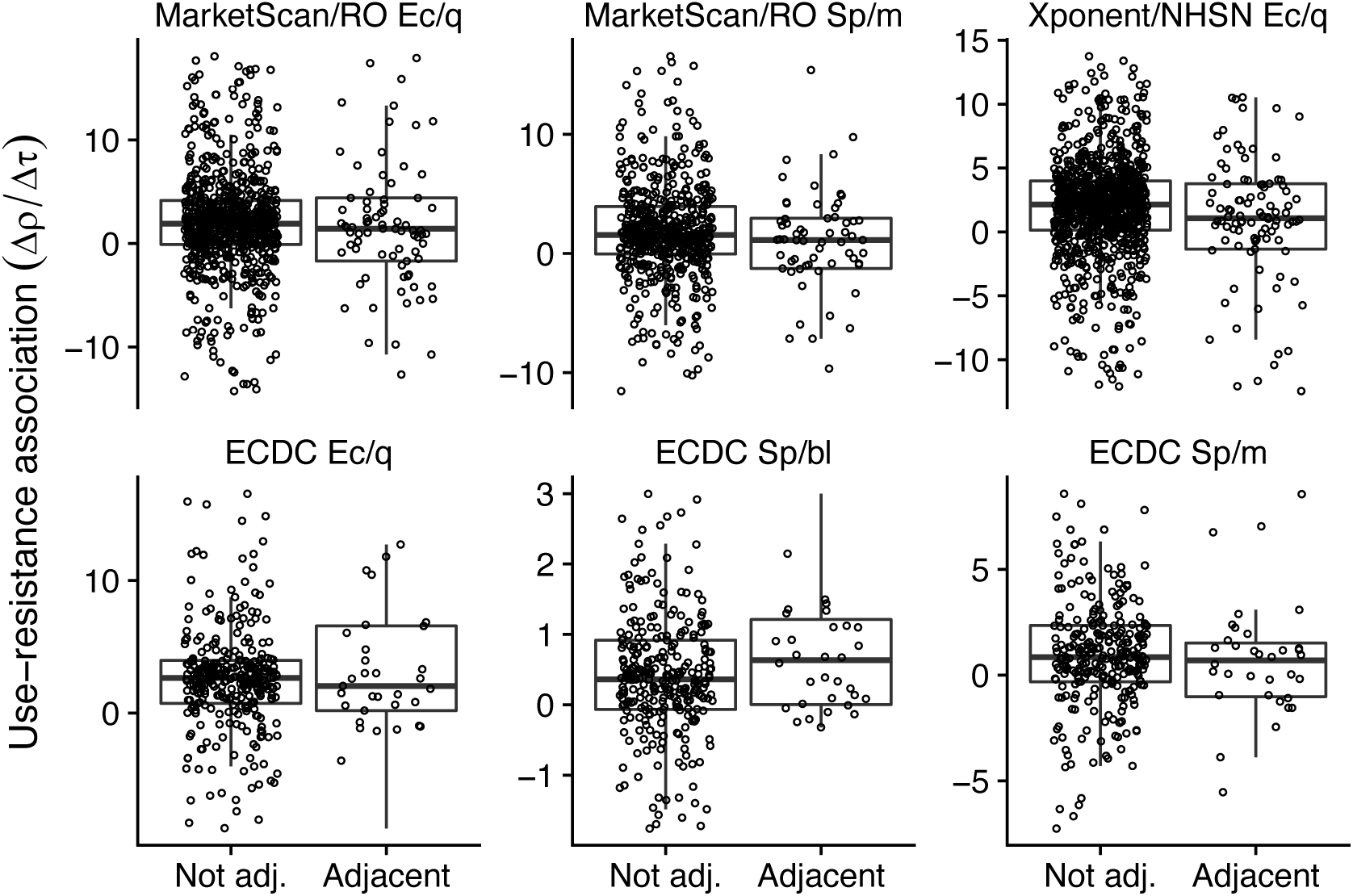
Use-resistance relationships by adjacency. Each point represents the use-resistance association in populations (top row, US states; bottom row, European countries), the same as those shown in Figure 2, arranged by whether the pair of populations is physically adjacent. Physically adjacent populations tend to have weaker use-resistance associations, but differences were not statistically significant. For visual clarity, the vertical axes are truncated to show only the central 90% of data points. Ec/q: *E. coli* and quinolones. Sp/m: *S. pneunomiae* and macrolides. Sp/bl: *S. pneumoniae* and β-lactams. RO: ResistanceOpen. ECDC: European CDC.

Next, to account for the possibility that adjacency was too coarse a measure for interactions between populations, we instead used a rank-ordered estimate of interactions using US commuting and European airline passenger flows (Supplemental Figure 2). In 4 of 6 dataset/pathogen/antibiotic combinations, the nonparametric association between increased inter-population interactions and decreased use-resistance associations was statistically significant, thus confirming the general trend observed in the adjacency analysis (Figure 4, Supplemental Table 3). The weakest significant result was for *S. pneumoniae* and macrolides in the MarketScan/ResistanceOpen dataset (Spearman’s *ρ* 0.07, 95% jackknife confidence interval -0.03 to 0.18; *p* = 0.028, Mantel test), and the strongest was for *E. coli* and quinolones in the Xponent/NHSN dataset (*ρ* = 0.13, 95% jackknife confidence interval 0.007 to 0.25; *p* = 0.001). The correlation for *S. pneumoniae* and macrolides in the ECDC data has a point estimate suggesting spillover but was not statistically significant, while the correlation for *S. pneumoniae* and β-lactams in the ECDC data had the opposite point estimate, consistent with the adjacency analysis (Figure 3). Finally, to quantify the effect of increased interactions on the observed use-resistance associations, we compared the use-resistance associations in pairs of populations within the lowest decile of interactions against those in the highest decile, using the same approach as for the adjacency analysis above (Supplemental Table 4). In 5 of 6 dataset/pathogen/antibiotic combinations, the point estimate for the different in use-resistance associations was consistent with spillover, with a weaker association among pairs of populations with greater interactions. In those 5 cases, the point estimates ranged from a 18% reduction up to a 75% reduction in use-resistance associations among the highest-interacting pairs of populations, compared to the lowest-interacting populations.

**Figure 4.**
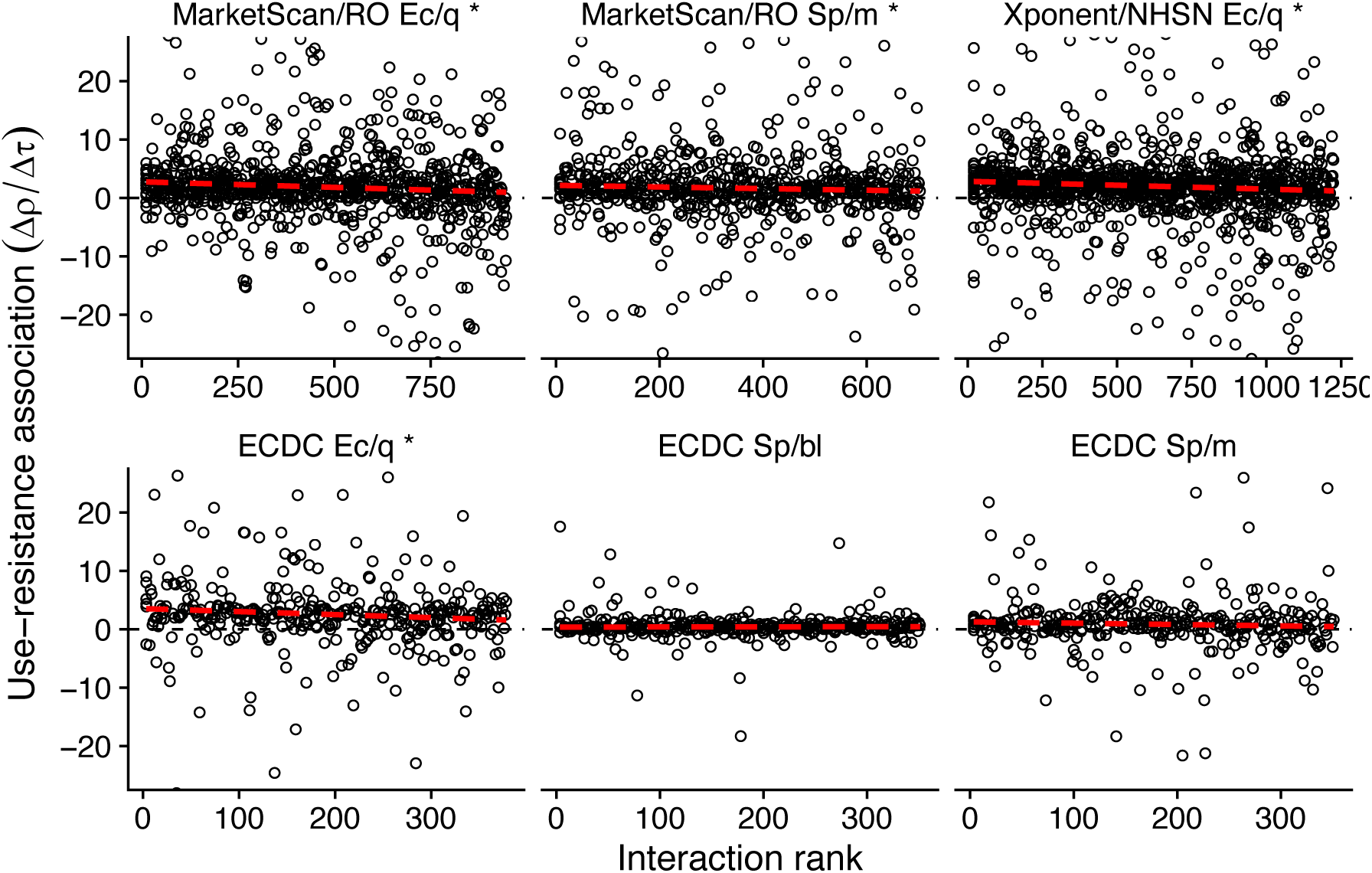
Use-resistance associations by ranked interaction. Each point represents the use-resistance association in a pair of US states (top row) or European countries (bottom row), the same pairs as shown in Figure 3, rank ordered by increasing inter-population interaction as inferred from transportation data. For visual clarity, the horizontal axes are truncated to exclude outliers. The dashed red line is a visual illustration of how increasing interaction is correlated with decreasing use-resistance associations (robust regression; compare Supplemental Tables 3 and 4). The asterisk (*) indicates a statistically significant association between increased interaction and decreased use-resistance relationship (Supplemental Table 3). Ec/q: *E. coli* and quinolones. Sp/m: *S. pneunomiae* and macrolides. Sp/bl: *S. pneumoniae* and β-lactams. RO: ResistanceOpen. ECDC: European CDC.

## DISCUSSION

We used theoretical models to show that interactions between two populations can attenuate the observed use-resistance association. In simulations, the quantitative relationship between inter-population interactions and the attenuation of the use-resistance association was dependent on the theoretical model used. However, we found that, in two models of the use-resistance association, having on the order of 1% of interactions between a control and intervention population was sufficient to attenuate the observed effect of theoretical stewardship intervention by 50%, relative to a situation where the two populations were completely isolated. These theoretical results suggest that even small numbers of interactions could lead to substantial spillover.

When examining observational antibiotic use and resistance data from US states and European countries, we did not detect a robust signal of spillover among pairs of adjacent populations, as opposed to non-adjacent pairs, even across 3 pathogen-antibiotic combinations in 3 separate datasets. However, when using more fine-grained transportation data to estimate the relative ranking of epidemiological contacts between those populations, we found a correlation between increased interactions and attenuated use-resistance associations. Pairs of populations in the highest decile of inter-population interactions, that is, those most subject to spillover, had use-resistance associations on the order of 50% weaker than pairs in the lowest decile of interactions. The 2 pathogen/antibiotic dataset combinations with data not indicative of spillover, namely *S. pneumoniae* and β-lactams and macrolides in the ECDC data, may have not shown the same signal as other cases because of the smaller number of populations in those cases (27, versus 28 to 50 in the other cases) led to insufficient statistical power or potentially because the biology or epidemiology of *S. pneumoniae* resistance in these cases is somehow different and does not exhibit spillover.

These theoretical and empirical results suggest that spillover is relevant at the level of US states and European countries. This finding has important ramifications. First, attempts to attribute changes in a population’s level of antibiotic resistance to changes in that population’s rates of antibiotic use may lead to inaccurate conclusions unless use and resistance in surrounding populations is accounted for. Second, state-or country-level antibiotic stewardship pilot studies may substantially underestimate the potential reduction in antibiotic resistance that would follow from a reduction in antibiotic use if that reduction were implemented at a larger scale. Third, mass drug administration trials may lead to elevated levels of antibiotic resistance in the control populations if those populations are not entirely separated from the intervention population. Finally, spillover can at least partly explain why use-resistance associations at the level of US states or European countries are sometimes difficult to detect and, when they are detected, are sometimes weaker than expected (5,11,14). Furthermore, spillover means that theoretical models of antibiotic use and resistance that treat US states or European countries as epidemiologically independent populations will not accurately represent the dynamics of resistance (33).

Our study has several limitations. First, we interpreted the theoretical results and ecological data as if the association between antibiotic use and resistance were causal and deterministic. However, decreases in the use of an antibiotic may not necessarily lead to declines in resistance to that antibiotic in a target pathogen (12,47–49). We do not address co-resistance and cross-selection (50,51), and we assumed that resistance equilibrates on a timescale comparable to an intervention. Previous research has shown that resistance among *E. coli, S. pneumoniae, N. gonorrhoeae* and other organisms can respond to changes in antibiotic use on the timescale of months (52–55), but the expected delay between a perturbation to antibiotic use and the resulting change in resistance remains a subject of active study (13,52,56,57). Nevertheless, the use of ecological data was essential to addressing our hypothesis, as data from multiple controlled, state-or country-wide experiments are not available.

Second, our analyses attributed all differences in antibiotic resistance between populations to differences in use across those populations and to interactions between them. In fact, antibiotic resistance is associated with factors beyond antibiotic use (6,58), and those factor are likely spatially correlated. In other words, closely interacting populations might have more similar use-resistance associations because they tend to be more similar with respect to other determinants of antibiotic use. Our estimates of the correlation between inter-population interactions and the attenuation of use-resistance relationships may therefore be overestimates. A more careful quantification of the relative roles of spillover versus other spatially-correlated determinants of resistance is required.

Third, our analysis only considered pairs of populations, when in fact spillover is happening between all pairs of populations in our analysis simultaneously. We used the pairs approach because it allowed for a simple theoretical model and a straightforward comparison of theory with the observational data. However, more sophisticated approaches that account for the network of spillover interactions will likely lead to more refined characterizations of spillover.

Finally, analyses based on administrative entities like US states or European countries, although logistically attractive “laboratories” of antibiotic stewardship, will always be difficult to interpret because administrative entities average over important dimensions of population structure like age (59), sexual networks (60), and race/ethnicity (61). Thus, use-resistance associations measured across states and countries may be different from those that appear among geographically-proximate populations with dissimilar antibiotic use rates, such as the sexes (62) and racial/ethnic groups (63).

We suggest 3 lines of investigation that could refine our understanding about the role of spillover at levels of US states and European countries. First, further mathematical modeling studies with more realistic structuring of the host population might articulate more detailed theoretical expectations about the relationship between intervention scale and spillover. For example, models could be parameterized with epidemiological information about individuals’ contacts and travel patterns, as has been done for other infectious diseases (64). Second, meta-analysis of existing studies of use-resistance relationships (5,65,66), both experimental and observational, could potentially determine the empirical relationship between intervention population size and the importance of spillover. This kind of meta-analysis might reveal that populations other than US states are feasible “laboratories” for resistance: it may be that cities, daycares, schools, workplaces, or even families represent the optimal trade-off between maximizing logistical feasibility and minimizing spillover. Finally, future experimental outpatient antibiotic stewardship interventions should make careful and deliberate decisions about the sizes and interconnectedness of the populations they target. We hope that a better understanding of spillover will improve predictions about the future of antibiotic resistance, the formulation of stewardship policy, the design of stewardship interventions and antibiotic administration trials, and theoretical models of resistance.

## Disclaimers

The views and opinions of the authors expressed herein do not necessarily state or reflect those of the ECDC. The accuracy of the authors’ statistical analysis and the findings they report are not the responsibility of ECDC. ECDC is not responsible for conclusions or opinions drawn from the data provided. ECDC is not responsible for the correctness of the data and for data management, data merging and data collation after provision of the data. ECDC shall not be held liable for improper or incorrect use of the data.

## Acknowledgements

We thank Dr. Stephen M. Kissler for helpful comments on the manuscript.

