## Supplemental Information for "The role of “spillover” in antibiotic resistance"

### Supplemental Methods

*WHN model*

In the WHN model, there are *N* hosts who may be uncolonized *X*, carry a susceptible strain *S*, carry a resistant strain *R*, or be dual carriers *D*. Dual carriers are predominantly sensitive-colonized and do not transmit the resistant strain. Carriers naturally clear at a rate *u*. Antibiotics, used by all hosts at a rate *τ*, clear susceptible organisms. A fitness cost *c* reduces the resistant strain's transmission rate.

$$
\begin{aligned}
\dot{S} &= \beta \frac{S+D}{N} X - (u+\tau)S - \beta(1-c)\frac{R}{N}S \\
\dot{R} &= \beta(1-c)\frac{R}{N}X - uR + \tau D \\
\dot{D} &= \beta(1-c)\frac{R}{N}S - (u+\tau)D \\
\dot{X} &= -(\dot{S}+\dot{R}+\dot{D}) \\
N &= X+S+R+D
\end{aligned}
$$

As per Davies *et al*. (1), we define the proportion of carriers carrying a resistant organism as $\rho = R / (S + R + D)$.

We use the parameterization in Davies *et al.*, designed to match the relationship between rates of β-lactam use and resistance among *Streptococcus pneumoniae* across European countries: $\beta = 4$ per month, $u = 1$ per month, and $c \approx 0.168$ (such that $\rho = 0.5$ for 1.5 treatments per year). We fixed the relative efficiency of multiple colonization (denoted *k* in the original presentation) at 1 in the equations above and in all simulations.

Simulations were initiated with *S* and *R* populations each accounting for 5% of hosts and *X* accounting for 90%. Simulations were run to equilibrium using the *runsteady* function in the *rootSolve* package (2) in R (version 3.6.0) (3).

*WHN model with 2 populations*

The transmission constant between populations depends on the interaction parameter $\varepsilon \in \left[0, 1 - \frac{1}{n}\right]$:

$$\beta_{ij} = \beta \times \begin{cases} 1 - \varepsilon & i = j \\ \varepsilon & i \neq j \end{cases}$$

The dynamical model is then

$$\dot{S}_i = \sum_j \beta_{ij} \frac{S_j + D_j}{N_j} X_i - (u + \tau_i) S_i - (1 - c) \sum_j \beta_{ij} \frac{R_j}{N_j} S_i$$

$$\dot{R}_i = (1 - c) \sum_j \beta_{ij} \frac{R_j}{N_j} X_i - u R_i + \tau_i D_i$$

$$\dot{D}_i = (1 - c) \sum_j \beta_{ij} \frac{R_j}{N_j} S_i - (u + \tau_i) D_i$$

$$\dot{X}_i = -\left(\dot{S}_i + \dot{R}_i + \dot{D}_i\right)$$

$$N_i = X_i + S_i + R_i + D_i$$

In our simulations, we set the population sizes $N_i$ equal.

*D-types model*

In the D-types model (4), there are $n_D$ “duration”-types, each with clearance rate $u_d$. Each type *d* has sensitive and resistant strains, so the compartments are uncolonized *X*, sensitive-colonized $S_d$, and resistant-colonized $R_d$. The D-types are subject to balancing selection encoded by

$$v_d = \left(1 - \left[\frac{S_d + R_d}{\sum_{d=1}^{n_D}(S_d + R_d)} - \frac{1}{n_D}\right]\right)^k$$

where $k \geq 1$ is the force of that selection. Thus:

$$\begin{aligned} \dot{S}_d &= v_d \beta \frac{S_d}{N} X - (\tau + u_d) S_d \\ \dot{R}_d &= v_d \beta \frac{R_d}{N} X - c u_d R_d \end{aligned}$$

In this model, the cost of antibiotic resistance *c* is associated with increased clearance of resistant strains rather than reduced transmission. The proportion resistant is

$$\rho = \frac{\sum_d R_d}{\sum_d (S_d + R_d)}$$

We extend the single-population D-types model to a multi-population model analogously to the WHN model:

$$\begin{aligned} \dot{S}_{i,d} &= v_{i,d} \sum_j \beta_{ij} \frac{S_{j,d}}{N} X_i - (\tau_i + u_d) S_{i,d} \\ \dot{R}_{i,d} &= v_{i,d} \sum_j \beta_{ij} \frac{R_{j,d}}{N} X_i - c_\mu u_d R_{i,d} \end{aligned}$$

where $S_{i,d}$ is the number of hosts in population *i* carrying the susceptible variant of the type *d*.

The model parameters are drawn from the original publication: $n_D = 16$, *β* = 2 per month, *c* = 1.1, *k* = 15, and the $u_d$ evenly spaced from 0.5 to 2 per month.

*MarketScan/ResistanceOpen data*

State-level MarketScan and ResistanceOpen data with masked state labels were published previously (5). The masked labels are not linked with geographic information and so cannot be used to reproduce the results in this study. The ResistanceOpen data are mostly drawn from hospitals or health systems, using CLSI laboratory standards, and covering a variety of specimen types (6).

*Xponent/NHSN data*

Quinolone use data from the Xponent database was filtered for years between 2011 and 2014. State-level values used in the analysis were the average of the use rates over those 4 years. *E. coli* resistance data included all isolates from all age groups and all years 2011-2014.

*ECDC data*

Antibiotic use and resistance data were filtered for the years 2011-2015. Numbers of non-susceptible and total isolates were set as the sum of the yearly numbers. Antibiotic resistance data is drawn from blood and cerebrospinal isolates using the EUCAST laboratory methodology (7). Country-level use values were set as the average of the yearly use rates. Ambulatory care antibiotic use rates were used. When data was not available for some pathogen-antibiotic-country-year combinations, the country-level averages were taken over the remaining years. Exceptions are shown in the table below.

| Data | Exceptions to 2011-2015 time range |
|---|---|
| Quinolone, β-lactam, and macrolide use | Iceland (2014-2015), Lithuania and Slovakia (2012-2015), Poland (2014-2015), Sweden (2014-2015) |
| Macrolide resistance among *S. pneumoniae* | Latvia (2012-2015), Luxembourg (2014-2015) |

To facilitate comparison of European and US use data, the European use rates, measured in defined daily doses (DDD) per 1,000 inhabitants per day (DID), were converted to treatments per person per year by assuming that 10 DDD of β-lactams equal one treatment, 10 DDD of quinolones equal one treatment, and 7 DDD of macrolides equal one treatment (8). This conversion is only for visualization purposes.

*Code and data availability*

Code to reproduce the theoretical simulations, the Xponent/NHSN and ECDC data, and code to reproduce the observational analysis on those two datasets is online at DOI: 10.5281/zenodo.3909812. Xponent/NHSN data on *E. coli* and quinolones is publicly available from the US CDC website (9,10). ECDC data on the 3 pathogen-antibiotic combinations is publicly available from ECDC websites (11,12).

**Supplemental Figure 1.** The dependence of the use-resistance relationship ($\Delta\rho/\Delta\tau$, vertical axis) on the interaction strength $\varepsilon$ in the “D-types” model for two differences in antibiotic use $\Delta\tau$. Compare Figure 1e, which shows this result for the WHN model.

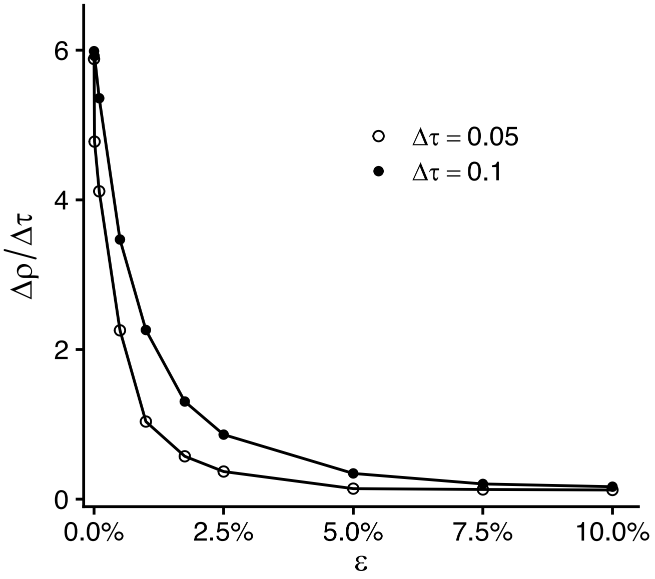

**Supplemental Figure 2.** Adjacency of US states and European countries according to ranked interaction deciles. US states with the greatest number of interactions (measured by commuting flows) are mostly adjacent, while adjacent European countries do not tend to have stronger interactions (as measured by passenger flights).

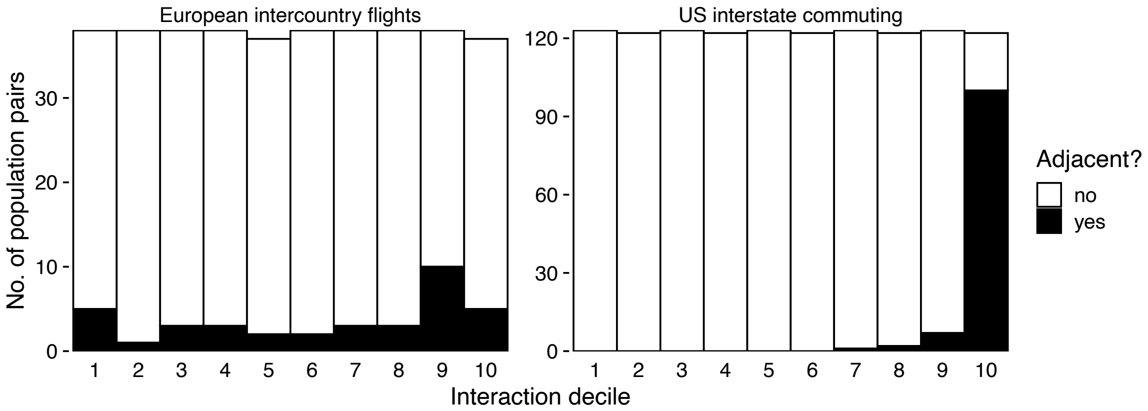

**Supplemental Table 1.** Reductions in the use-resistance relationship ($\Delta\rho/\Delta\tau$) depending on inter-population interactions $\varepsilon$, relative to the case with no interaction ($\varepsilon$ = 0). Compare Figure 1e and Supplemental Figure 1.

| | **Reduction in $\Delta\rho/\Delta\tau$** | | | |
|---|---|---|---|---|
| | **WHN model** | | **"D-types" model** | |
| **$\varepsilon$** | **$\Delta\tau$ = 0.05** | **$\Delta\tau$ = 0.10** | **$\Delta\tau$ = 0.05** | **$\Delta\tau$ = 0.10** |
| 0% | 0% | 0% | 0% | 0% |
| 0.01% | 0.4% | 0.7% | 19% | 1% |
| 0.1% | 4% | 6% | 30% | 11% |
| 1% | 29% | 32% | 82% | 62% |
| 10% | 41% | 43% | 90% | 78% |

**Supplemental Table 2.** The difference in median use-resistance association ($\Delta \rho/\Delta \tau$) between the adjacent and non-adjacent pairs (negative numbers signify that adjacent pairs have smaller $\Delta \rho/\Delta \tau$) and the corresponding fractional change. For example, a fractional change of -0.26 means a reduction of 26%. Compare Figure 2. Intervals are jackknife 95% confidence intervals.

| **Dataset** | **Difference in median $\Delta \rho/\Delta \tau$** | **Fractional change in median $\Delta \rho/\Delta \tau$** | ***p* value (Mann-Whitney *U*)** |
|---|---|---|---|
| MarketScan/RO Ec/q | -0.49 (-1.7 to 0.67) | -0.26 (-0.82 to 0.31) | 0.48 |
| MarketScan/RO Sp/m | -0.42 (-1.2 to 0.38) | -0.27 (-0.7 to 0.16) | 0.098 |
| Xponent/NHSN Ec/q | -1.1 (-2.2 to 0.053) | -0.50 (-0.97 to -0.038) | 0.020 |
| ECDC Ec/q | -0.61 (-4.1 to 2.9) | -0.23 (-1.5 to 1.1) | 0.97 |
| ECDC Sp/bl | 0.27 (-0.49 to 1.0) | 0.75 (-1.5 to 2.9) | 0.24 |
| ECDC Sp/m | -0.15 (-2.4 to 2.1) | -0.18 (-2.7 to 2.4) | 0.25 |

**Supplemental Table 3.** Correlations (Spearman's $\rho$) between a population pair's use-resistance association $\Delta\rho/\Delta\tau$ and ranked interaction. Compare Figure 4. Intervals are jackknife 95% confidence intervals.

| **Dataset** | **Correlation** | ***p* value (Mantel test)** |
|---|---|---|
| MarketScan/RO Ec/q | -0.11 (-0.22 to -0.0032) | 0.002* |
| MarketScan/RO Sp/m | -0.072 (-0.18 to 0.033) | 0.029* |
| Xponent/NHSN Ec/q | -0.13 (-0.25 to -0.007) | 0.001* |
| ECDC Ec/q | -0.15 (-0.34 to 0.046) | 0.004* |
| ECDC Sp/bl | 0.022 (-0.19 to 0.24) | 0.66 |
| ECDC Sp/m | -0.086 (-0.29 to 0.12) | 0.072 |

* Statistically significant after multiple hypothesis correction.

**Supplemental Table 4.** Absolute and fractional difference in use-resistance association $\Delta\rho/\Delta\tau$ among pairs of populations in the top decile of interactions compared to pairs in the bottom decile. Negative differences indicate a spillover effect: more interactions correlate with weaker use-resistance relationships. Negative ratios also show a spillover effect. For example, a fractional change of -0.54 indicates a 54% reduction in median use-resistance relationships in the highest-interacting decile pairs of populations compared to the lowest-interacting decile pairs. Compare Figure 4. Intervals are jackknife 95% confidence intervals.

| **Dataset** | **Difference in $\Delta\rho/\Delta\tau$** | **Fractional change in $\Delta\rho/\Delta\tau$** |
|---|---|---|
| MarketScan/RO Ec/q | -1.1 (-2.3 to 0.20) | -0.54 (-1.1 to -0.017) |
| MarketScan/RO Sp/m | -0.63 (-2.7 to 1.5) | -0.38 (-1.5 to 0.77) |
| Xponent/NHSN Ec/q | -1.6 (-2.7 to -0.40) | -0.58 (-0.95 to -0.21) |
| ECDC Ec/q | -0.55 (-4.3 to 3.2) | -0.18 (-1.2 to 0.79) |
| ECDC Sp/bl | 0.18 (-0.71 to 1.1) | 0.51 (-7.8 to 8.8) |
| ECDC Sp/m | -0.64 (-1.6 to 0.33) | -0.75 (-1.9 to 0.35) |
